# Hunting for microsatellite instability in long-read data with Owl

**DOI:** 10.64898/2026.03.08.708905

**Authors:** Zev Kronenberg, Khi Pin Chua, Mark J.P. Chaisson, Byunggil Yoo, Lisa Lansdon, William J. Rowell, Guilherme de Sena Brandine, Egor Dolzhenko, Kobe Ikegami, Kie Kyon Huang, Patrick Tan, Shruti Bhise, Everett Fan, Mark Mendoza, Emily O’Donnell, Tomi Pastinen, Elizabeth R. Lawlor, Scott N. Furlan, Midhat S. Farooqi, Michael A. Eberle

## Abstract

Microsatellite instability (MSI) is a key biomarker of mismatch repair deficiency and response to immunotherapy, yet most existing genomic detection methods are optimized for short-read sequencing and rely on small panels of homopolymer markers, limiting the ability to characterize genome-wide and motif-specific patterns of instability. Here we present Owl, a bioinformatic tool for quantifying MSI from long-read (PacBio) genomic data. Owl leverages a genome-wide marker set of more than 140,000 microsatellite repeats ranging from 1–6 bp in length to measure MSI across a phased genome. Using a wrap-around alignment algorithm, Owl constructs repeat-length distributions at each marker site and flags somatic instability using the coefficient of variation. We applied Owl to screen for markers with stable coverage, phasing, and baseline variation across 131 diverse genomes from the Human Pangenome Reference Consortium, where Owl scores ranged from 1.4% to 5.4% of markers exceeding the instability threshold. When applied to 19 cancer cell lines and one diffuse astrocytoma tumor-normal pair, Owl identified five MSI-high genomes with 15-18% unstable markers and showed close concordance with an Illumina DRAGEN MSI assay for the astrocytoma sample. Motif-level analyses revealed shared enrichment of short homopolymer and dinucleotide (A– and AT-rich) repeats across MSI-high cancers, and additionally uncovered a distinct pattern of elevated GGAA microsatellite instability in Ewing sarcoma cell lines, consistent with the known role of the EWS::FLI1 fusion protein at GGAA-rich regulatory elements. Owl is implemented in Rust and integrated into the PacBio HiFi Somatic workflow, providing a scalable framework for MSI analysis from long-read sequencing focused on repeat instability specifically in tumor samples.

## Introduction

Deficiencies in DNA repair pathways give rise to somatic mutations ranging from small variants (SNVs and indels) (Alexandrov et al. 2013) to large-scale structural variants (Sohn et al. 2025) that collectively drive cancer progression. Tracking deficiencies in the DNA repair pathway is typically done indirectly through several genomic metrics. The first is Tumor Mutational Burden (TMB), which measures the accumulation of somatic Single Nucleotide Variant (SNVs) and small insertions and deletions (indels). The second, Homologous Recombination Deficiency (HRD), is a compound metric that tracks the loss of heterozygosity, large-scale state transitions, and telomeric imbalance. Lastly, Microsatellite Instability (MSI) is when somatic insertions and deletions accumulate in simple repeats. Although these measures of genomic instability and scarring do not directly pinpoint the causative DNA repair defect, they remain critical biomarkers for cancer diagnosis and treatment planning (Gatalica et al. 2016; Telli et al. 2016; Choucair et al. 2020).

MSI-high cancers typically arise through loss of function mutations in one of several key Mismatch Repair (MMR) genes, including *MLH1*, *MSH2*, *MSH6*, and *PMS2* (Vilar and Gruber 2010). Lynch syndrome, for example, is an autosomal dominant hereditary cancer predisposition caused by germline mutations in the MMR genes, commonly *MLH1* and *MSH2* (Bhattacharya et al. 2025). In contrast, sporadic (non-syndromic) MSI-high cancers acquire somatic inactivation of these same repair genes in a classical “two-hit” manner. Beyond direct mutations, epigenetic silencing of *MLH1* through promoter hypermethylation is a well-established mechanism of MMR deficiency and a major driver of tumorigenesis in sporadic MSI-high colorectal and endometrial cancers (Niv 2007). MMR driven cancers (including colorectal, endometrial, gastric and ovarian cancers) frequently exhibit high MSI (Kavun et al. 2023). Although less common, MSI-high tumors have also been reported in prostate, pancreatic, and brain cancers (Bonneville et al. 2017).

Detection methods for microsatellite instability (MSI) from the genome include PCR, Sanger sequencing, and short-read sequencing (SRS) approaches (Yakushina et al. 2023). Genomic assays typically target loci containing simple repeats, such as the BAT-25 and BAT-26 markers (Suraweera et al. 2002), which are homopolymer tracts located within introns of *KIT* and *MSH2*. MSI is typically defined by >30% of classical PCR markers loci showing elevated rates of variability (Sahin et al. 2019). Although efforts to standardize biomarker selection for high-throughput sequencing are increasing (Andrews et al. 2024), existing frameworks such as the Bethesda guidelines (Boland et al. 1998) and their associated marker panels were developed for PCR-based assays and have not scaled to whole-genome sequencing (WGS). Ideal WGS-compatible markers should be stable across human populations, robust to technical artifacts, easily profiled using whole-genome (WGS) or targeted exome sequencing (ES), and particularly sensitive to replication errors characteristic of mismatch repair (MMR)-deficient cells.

The MSI status of tumors is important to determine the immunotherapies most likely to be effective (Kavun et al. 2023). For example, frameshift mutations can generate novel peptides, or neoantigens, that are recognized as foreign by the immune system and tumor-infiltrating T cells can detect these neoantigens and mount an immune response. However, this immune response is often suppressed by immune checkpoint pathways such as the PD-1/PD-L1 axis. Checkpoint inhibitor antibodies, including those targeting PD-1, release this inhibition and allow T cells to mount a stronger antitumor response. In patients with MSI-high tumors, these therapies have been associated with markedly improved clinical outcomes. Sahin and colleagues provide a comprehensive review of MSI-targeted immunotherapy (Sahin et al. 2019).

There have been a number of bioinformatic tools developed to detect MSI in short-read sequencing (SRS) data, encompassing both genome (GS) and exome (ES) sequencing. Modern SRS-based assays typically classify tumors as MSI-high when approximately 10-20% of evaluated loci are unstable, with the exact cutoff varying by algorithm and calibration dataset. Commonly used tools include MSIsensor (Niu et al. 2014), *MANTIS (Kautto et al. 2017)*, *MSIsensor-pro* (Jia et al. 2020), *mSING* (Salipante et al. 2014), and DRAGEN, most of which have been independently benchmarked by Anthony and Seoighe (Anthony and Seoighe 2024), who found that these methods generally perform better on ES data than on GS data. A key challenge in detecting MSI from short-read sequencing (SRS) data is distinguishing somatic mutations from germline heterozygosity, since SRS data are not phased. To address this limitation, some tools compare repeat-length distributions between paired tumor and normal samples. However, matched normal samples are not always available and add to assay cost, and SRS inherently struggles to resolve long or complex repeats due to mapping ambiguities.

Long-read sequencing (LRS), including PacBio HiFi and Oxford Nanopore Technologies, offers several advantages for MSI detection compared to SRS. Long reads span nearly all microsatellite regions, enabling accurate repeat localization through improved mapping, and long-read phasing provides haplotype-level resolution of repeat variation within microsatellites. However, LRS has historically been limited by higher indel error rates and greater cost relative to SRS. As LRS becomes more practical (i.e., higher throughput and lower sequencing cost), it is being adopted for tumor analysis of structural variation, direct profiling of DNA methylation, and full length transcript sequencing (Ermini and Driguez 2024; Li et al. 2025; Lansdon et al. 2024). A complete analysis of tumors sequenced with long reads requires software tools developed specifically to work with this data type.

In this study, we present *Owl* (**Figure 1A**), a bioinformatic tool for determining whether a sample exhibits MSI-high or microsatellite-stable (MSS) characteristics. *Owl* takes advantage of LR WGS, including precise repeat mapping and read-level phasing, to quantify the fraction of marker sites that exhibit MSI at haplotypic resolution. We examine variation at these marker sites across a diverse panel of human population samples (HPRC), cancer cell lines, and real-world tumor specimens to identify the most reliable indicators of MSI. Our results show that *Owl* provides highly accurate detection of MSI and establishes a robust framework for characterizing genomic instability in cancer.

**Figure 1.**
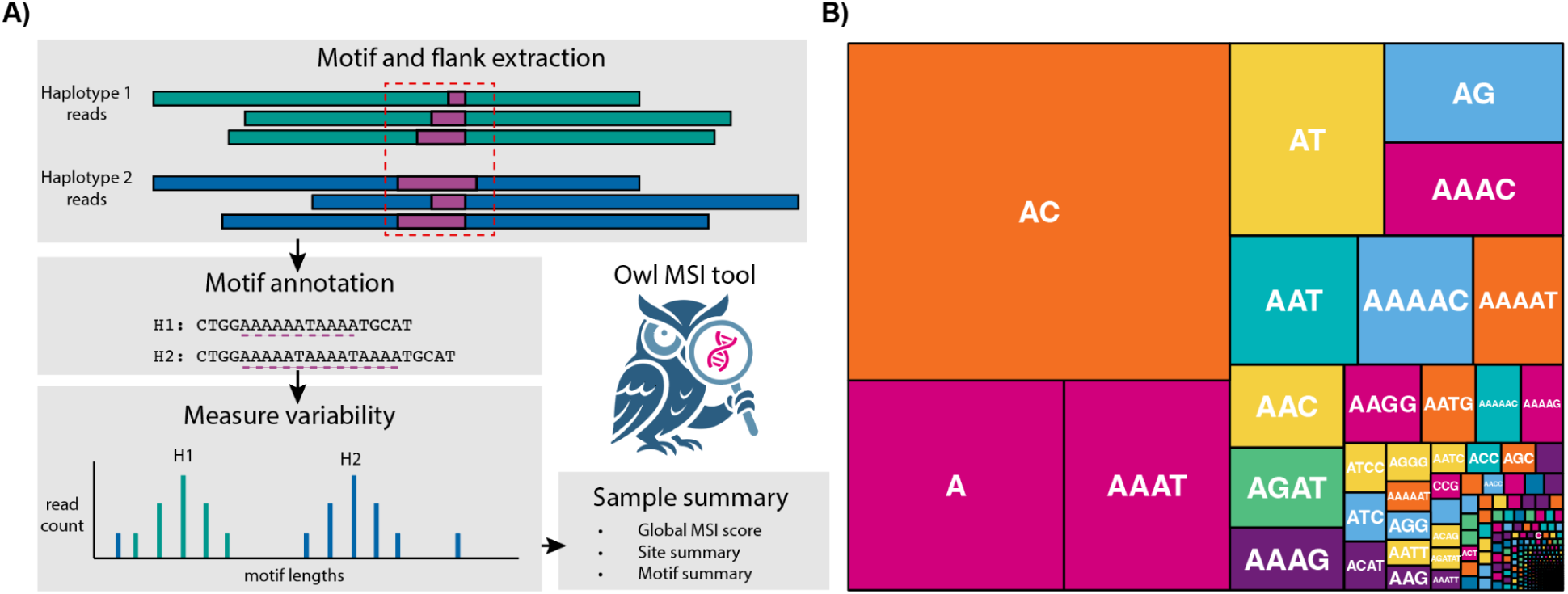
Owl workflow and marker set. **A**) The *Owl* workflow starts with a mapped and phased HiFi genome. In the first stage (profile), reads spanning predefined marker sites are extracted, including flanking bases. Repeats are then annotated in each read using a wrap-around dynamic programming algorithm to determine the matching motif repeat length. These repeat lengths are then grouped by haplotype, and for each marker, the mean, standard deviation, and coefficient of variation are calculated. In the second stage (*score*), *Owl* summarizes MSI across all markers and produces human-readable output files for both profiling and scoring. **B)** Treemap of repeat motif frequencies in the GRCh38 marker set used by Owl. Each box represents a unique (lexicographically minimized) repeat motif, with area proportional to its number of genomic sites. The plot shows motif sequences only; repeat lengths are not depicted.

## Results

*Owl* consists of two primary modules: profile, which processes read alignments; and score, which calculates and summarizes MSI at the sample level (**Figure 1A**). The profile module operates on aligned and phased HiFi reads together with a bed file of repeat regions and their corresponding motifs. For each marker, the repeat motif is annotated within reads using wrap-around alignment (**see Methods**). Reads are then grouped by haplotype, and for each locus, the coefficient of variation (CV) of repeat length is calculated, serving as the principal per-locus measure of instability used throughout *Owl*. The score module then aggregates the per-locus CV values genome-wide, reporting the percentage of markers that exceed the predefined CV threshold of five, derived from a control genome (**see Methods**). The output also includes summary statistics such as the fraction of phased markers, the proportion of loci exhibiting high CV, and a motif-specific breakdown of results.

We first identified a set of microsatellites to serve as the *Owl* MSI markers. Based on the GRCh38 RepeatMasker annotations from the UCSC Genome Browser (Kent et al. 2002), we selected simple 1-6 bp repeats spanning 20-100 bp with sequence identity greater than 97%. A high identity cutoff was required because the repeat annotation algorithm does not account for divergent or mixed repeats. Markers were restricted to canonical chromosomes (**Figure S1**) (excluding chromosome Y). This produced 164,374 repeat sites including 11.6% homopolymers, 45% dinucleotide repeats, and 8.6% trinucleotide repeats. There were 340 distinct motifs after lexicographically minimizing rotations and reverse complements of the original 1,573 motifs (**Table S1**). Notably, our identity filtering indirectly excluded most C/G homopolymers and di-nucleotides (48 remaining), which are depleted in the human genome due to its ∼60% AT bias. Methylation and subsequent deamination of C to T (Fryxell and Moon 2005) frequently disrupt poly-C tracts, leading to higher divergence among C (15%) and G (9%) homopolymers compared to A (5%) and T (7%).

For a marker/technology combination to best detect MSI, it is important that the repeats used do not show excessive signals of instability in control samples that may be caused by technical artifacts. We profiled all 164,374 markers in a set of control PacBio HiFi genomes derived from lymphoblastoid cell lines using *Owl* and the new marker panel (**Table S2**). The dataset included 131 diverse samples from the Human Pangenome Reference Consortium (HPRC) (Liao et al. 2023). From this analysis, we removed 17,790 markers that had a high no-call rate or CV leaving us with 146,562 high quality markers (**see Methods**). Across these controls, the *Owl* score, which is the percentage of markers classified as highly variable, ranged from 1.39% to 5.39%, with a mean of 2.18% and a standard deviation of 0.51 (**Figure 2A**). Weighted by total genomic counts, background instability varied by motif length, with 7.84% for homopolymers, 1.71% for dinucleotides, and 1.11% for trinucleotides. Although C or G homopolymers and CG dinucleotides showed high apparent instability (61% and 29%, respectively), these motifs are extremely rare in the marker set and together account for only 48 sites. Their low abundance makes the instability percentage highly sensitive to sampling noise. Among the 1,287 motifs with more than 10% high across any control sample, only 21 (1.6%) have more than 100 genomic sites per individual. This shows that motifs with high apparent instability are predominantly low-frequency, and their overall contribution to the MSI score is naturally minimized by their low genomic abundance.

**Figure 2.**
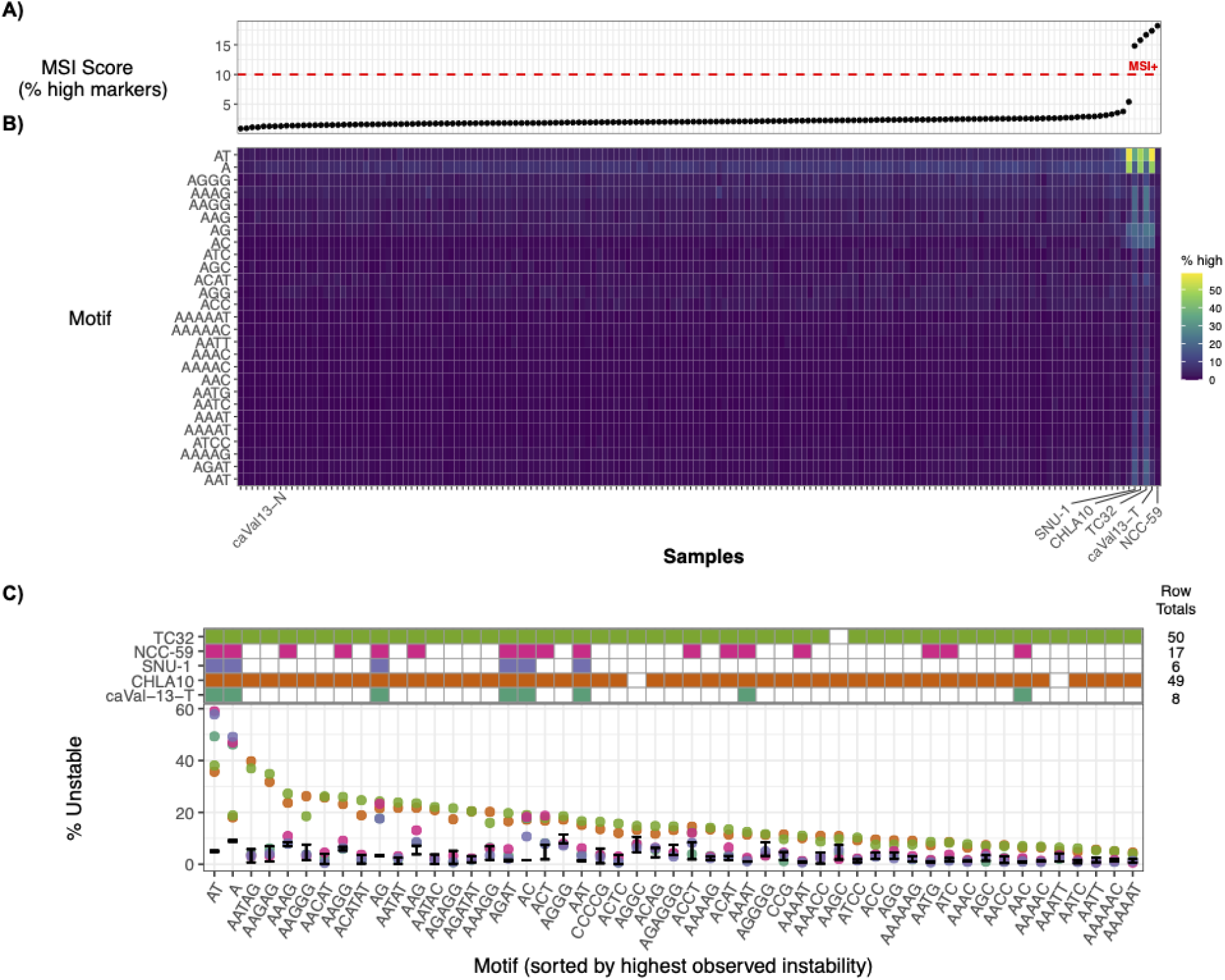
Per-sample MSI scores and motif-specific microsatellite variability. **A**) Owl MSI scores for each sample, sorted from low to high. Two Ewing sarcoma cell lines (CHLA10 and TC32), two gastric cancers (SNU-1 and NCC-59), and one diffuse astrocytoma (caVal13) exceed the 10% MSI threshold. **B)** Heatmap showing the percentage of high-variability sites per motif, restricted to the most frequent motifs (>500 marker loci). Samples along the x-axis follow the order from the panel above, while motifs along the y-axis are clustered by similarity using hclust. The five MSI-high samples are labeled and the diffuse astrocytoma control (N). **C)** Outlier motifs in the five MSI-high samples. The top panel marks which sample contains a Bonferroni-corrected p-value (p<0.05) for a beta-binomial one-sided test. The bottom panel shows, for each motif (lexicographically and rotationally minimized), the percentage of sites with high CV. Each point represents one of the five samples. The beta-binomial credible intervals (95%) for the HPRC samples are shown as black lines. Only motifs with >100 markers across the genome are shown.

Across the full dataset, we analyzed nine Ewing sarcoma cell lines (tumor only), one melanoma cell line with its matched control, four breast cancer cell lines with controls, two lung cancer cell lines with controls, two gastric carcinoma cell lines (tumor only), and one diffuse astrocytoma with its matched control (**Table S2**). Relative to the natural ∼1-6% range of instability observed in healthy genomes, five outlier cancer samples showed markedly elevated *Owl* scores ranging from 15-18% (**Figure 2A, Table S2**). These MSI-high samples included two Ewing sarcoma lines (TC32 and CHLA10), the diffuse astrocytoma (caVal13-T), and two gastric cancer samples (NCC-59 and SNU-1).

Next, we analyzed the baseline of motif specific instability across the 131 HPRC controls (**Figure 2B**). We modeled the controls with a hierarchical beta-binomial model that pools counts across samples for each motif and generates posterior-predictive p-values to flag motifs whose observed instability exceeds expectation (**see Methods**). Across the MSI-high samples, 80 motifs remained significant after Bonferroni correction (α = 0.05). However, only 51 of these were supported by more than 100 genomic markers. Six motifs (A, AT, AG, AC, AAT, AGAT) were found to be unstable in all five MSI-high samples (**Figure 2B & 2C**). The shared motifs suggest that the shorter motifs (1-3 bp) represent the biological signal of MSI. Using a single available tumor-normal pair, caVal-13, a diffuse astrocytoma, we found a negative correlation between motif length and coefficient of variation (CV) scores, with the strongest differences observed in homopolymers and dinucleotide repeats, and only modest changes in trinucleotide motifs (**Figure S4**). Among dinucleotides, the AT motif showed the largest average difference between tumor (5.68) and normal (2.65) CV scores. Although homopolymer and dinucleotide repeats often display elevated CVs even in microsatellite-stable samples (**Figure 2B**), the magnitude of variability was substantially higher in the cancer sample, providing a clear signal differentiation.

Interestingly, two of the nine Ewing sarcoma cell lines, TC32 and CHLA10, exhibited a distinct pattern of repeat instability (**Figure 2C**). Both showed a high proportion of significant motifs (∼98%; 50 and 49 motifs, respectively), notably more than the other MSI-high samples. A striking feature of this signal is its specificity to motifs close in edit distance to GGAA, a sequence bound by EWS::FLI1 at promoter and enhancer elements (Johnson et al. 2017). Whereas the other cancers were dominated by extreme A/AT outlier motifs, the Ewing lines displayed broadly elevated instability across GGAA-like motifs. In our analysis, TC32 and CHLA10 showed approximately threefold higher instability rates (23-26%) of GGAA motifs (lexicographically minimized to AAGG) scoring as unstable compared to other cancers and controls (3.1-9.7%; **Figure 2C**, **Figure S3**). This GGAA-specific pattern is particularly noteworthy because conventional MSI assays rely almost exclusively on homopolymers, meaning that Ewing sarcoma samples with elevated GGAA repeat instability may be classified as MSI-stable by standard tests, and it further suggests a mechanistic link between the observed microsatellite variation and the regulatory landscape shaped by EWS::FLI1 in Ewing sarcoma.

To further examine the relationship between GGAA repeat loci and transcriptional regulation, we first tested whether all GGAA-containing *Owl* markers were enriched within ENCODE (ENCODE Project Consortium et al. 2020) regulatory elements (**see Methods**). Of the 1,132 GGAA markers in our full set, 29% overlapped regulatory elements (p = 0.0005), consistent with prior reports that GGAA motifs are generally enriched in enhancer regions (Gangwal et al. 2008). We next assessed enrichment for all MSI-high *Owl* markers in the TC32 cell line, which showed a significant depletion (p = 0.0005). Finally, we evaluated the subset of GGAA MSI-high markers in TC32. Among the 520 MSI-high GGAA loci, 35% overlapped cis-regulatory elements, representing a significant enrichment (p = 0.0005). A similar pattern was observed in the CHLA10 cell line, where 36% of the 471 MSI-high GGAA markers intersected enhancer elements (p = 0.0005), whereas the total CHLA10 high-markers showed no enrichment (p=0.002).

Despite the background enrichment of GGAA motifs in regulatory regions, we identified two notable MSI-high GGAA loci that fall within genes relevant to Ewing sarcoma biology. First, we observed GGAA instability at a distal enhancer (EH38E2599658) within the *EZH2* locus. *EZH2* is transcriptionally upregulated by *EWS::FLI1* (Staege et al. 2004) and contributes to Ewing sarcoma tumorigenesis (Richter et al. 2009) (**Figure S5**). Second, we detected variable GGAA motifs that overlaps enhancers (EH38E1599035, EH38E1523911) within the gene bodies of *SOX5* and *SOX6* which has been shown to be activated by the binding of *EWS::FLI1* to a intronic GGAA-microstalite (Marchetto et al. 2020). In total, we found 151 genes that were within 1 kb of a GGAA instability overlapping an enhancer (**Table S5**). As additional long-read cancer genomes become available, we expect to gain a clearer understanding of how MSI-high GGAA-microsatalites impact the regulatory landscapes in Ewing sarcoma.

To evaluate how MSI calls derived from phased long-read data relate to those obtained from short-read sequencing, we compared the instability detected by Owl in HiFi whole-genome data with the instability detected by the DRAGEN MSI caller in matched exome short-read data from the same diffuse astrocytoma sample. DRAGEN identified 20% of its 12,315 evaluated homopolymer loci as unstable, while *Owl* detected 16.67% unstable sites across its long-read marker set (**Table S2**). When DRAGEN was run in tumor-only mode using a panel of normals as controls, the fraction of unstable loci decreased from 20% to 11%, illustrating how the underlying analysis mode influences MSI estimates in short-read assays. Despite differences in input data and marker sets, both approaches consistently classified the sample as MSI-high using lower, genome-wide thresholds (approximately 10-20%), reflecting the broader marker coverage relative to legacy PCR-based MSI assays (∼30%).

## Discussion

We present *Owl*, the first microsatellite instability (MSI) detection tool developed specifically for long-read sequencing data from cancer samples and optimized for PacBio HiFi data. Unlike short-read MSI callers, *Owl* leverages full-length repeat coverage and haplotype phasing to quantify variability in simple sequence repeats across the genome. *Owl* provides a genome-wide instability score that integrates directly into the PacBio HiFi somatic workflow (Chua 2025). *Owl* is concordant with DRAGEN at the genome-wide level, underscoring the robustness of long-read MSI profiling.

A key advantage of *Owl* is that it can operate using tumor-only samples. Because short-read methods do not phase reads, variation across haplotypes must be analyzed in aggregate, forcing reliance on either a matched normal or a panel of normals to distinguish true somatic microsatellite mutations from pooled haplotypic variation. Likewise, when *Owl* is forced to collapse haplotypes across all markers, it shows substantially inflated repeat-length variation (∼28%) compared with the ∼1% variation observed when phase information is leveraged. However, by preserving haplotype structure, *Owl* can accurately resolve somatic microsatellite changes without matched controls, an advantage that is particularly important in settings where obtaining normal tissue is challenging or not feasible.

*Owl* is also designed to capture motif-specific patterns of microsatellite instability, enabling a more granular view of repeat variation than a single genome-wide score. By comparing each motif’s instability against background rates derived from 131 HPRC control genomes, we found that homopolymer and dinucleotide repeats show the strongest enrichment of instability across all MSI-high samples, making them the most sensitive reporters of variation in long-read data. Our analysis also uncovered a distinct signature in Ewing sarcoma cell lines, where we found that GGAA motifs and motifs at close edit distance to GGAA displayed markedly elevated instability and frequent overlap with enhancer elements. These findings demonstrate that motif-level MSI profiling with long reads can reveal both shared and cancer-specific patterns, including classes of unstable motifs that are largely invisible to standard assays focused only on homopolymers.

The present analyses are limited by the number of long-read cancer genomes currently available, and the comparisons presented here are restricted to a single tumor sample. As additional tumor types are sequenced, we anticipate that *Owl* will enable the discovery of other cancer-specific MSI patterns and motif-level signatures that may reflect distinct biological mechanisms. Future development will focus on improved per-locus specificity, expanding support for targeted long-read panels to enable higher-throughput applications, and extending the framework to RNA to explore instability signatures at the transcript level. As new methods for somatic phasing become available, these capabilities could also be integrated to better resolve allele-specific instability within tumor genomes.

Together, these results establish *Owl* as a robust framework for MSI detection from long-read data, extending the analytical reach of HiFi sequencing in cancer genomics and providing a foundation for future studies of repeat instability across diverse tumor types.

## Methods

### Owl MSI detection workflow

The *Owl* microsatellite instability detection tool was implemented in Rust and provides three subtools: profile, merge, and score, which are described below. The software is public and distributed under [PacBio’s license](https://github.com/PacificBiosciences/owl/blob/main/LICENSE.md) on github: https://github.com/PacificBiosciences/owl. Both the source code and pre-compiled static binaries are available. *Owl* has two inputs, an aligned and haplotagged bam file, and a motif bedfile containing regions and their motifs, see provided example in the repository (https://github.com/PacificBiosciences/owl/tree/main/data). Users may generate their own marker files or reproduce the provided file (https://github.com/PacificBiosciences/owl/analysis/MARKERS.md).

During the profiling phase, *Owl* queries the BAM files for genomic regions corresponding to each repeat marker. Alignments fully spanning these regions are filtered and processed. Owl should be run on reads that have already been haplotagged by tools like Hiphase (Holt et al. 2024) or WhatsHap (Patterson et al. 2015). For each read, the portion of the read overlapping a tandem repeat is determined by the base-level alignment, and is extracted along with a flanking sequence (10 bp, total) for context.

Local wrap-around dynamic programming (WDP) (Landau et al. 2001; Myers and Miller 1989) is used to score an alignment of an optimal number of copies of a motif. Briefly, WDP finds the optimal local alignment of a sequence S to T*, the Kleene closure of the motif sequence T (e.g. {NULL, AT, ATAT, ATATAT…} for motif AT). The dynamic programming recursions are defined for regular expression matching (Myers and Miller 1989; Gusfield 2010). The scoring matrices are defined to exclude alignments with fewer than 50% matches. Only the most frequently occurring motif is considered, in contrast to the string decomposition approach used for annotating motif diversity of centromere (Dvorkina et al. 2020) and VNTR sequences (Ren et al. 2023) where motif libraries are considered. The function returns an alignment score and the coordinates of the repeat span within the read, allowing each read to be assigned a single repeat length for that marker. Reads with low alignment scores (<90% identity) are excluded from further analysis.

After annotation, repeat lengths are grouped by their haplotype tag (HP). The mean and standard deviation of repeat lengths are then calculated for each haplotype to summarize copy variation within the sample. While the standard deviation should be sufficient to describe repeat variability, there is a positive relationship with the length of the repeat and the standard deviation. In other words, large repeats tend to have larger outliers (with respect to length), and increased standard deviation. This often happens when a few read alignments place the repeat insertion outside the target interval. To account for this issue, rather than filtering outliers, the standard deviation of repeat length is normalized by the mean repeat length. The coefficient of variation (CV), accounts for larger means having higher variability and allows better comparison across datasets. At the end of the Owl profile stage, the program reports the site, motif, and, for each haplotype, the coverage, mean repeat length, coefficient of variation expressed as a percentage (CV), and repeat lengths (when using the –keep-lengths option). The output is a text file modeled after the Variant Call Format (VCF), with a header that defines each field and a structure that supports merging multiple samples into a single file.

To determine an empirical cutoff for the coefficient of variation (CV), the metric used to quantify marker instability, we fitted parametric distributions to a control cell line (NA12878)(Kronenberg et al. 2025). Using the *fitdistrplus* [R] package (Delignette-Muller and Dutang 2015), normal, gamma, and Weibull distributions were evaluated. Sites with a CV of zero (14%) were excluded, as invariant loci do not capture sequencing error and would require a mixture of models to represent accurately. The upper tail of the distribution was also trimmed at the 99.9th percentile (CV = 20), as extreme values likely reflect alignment artifacts, phasing errors, or incorrect repeat unit identification. Among the models tested, the gamma distribution provided the best fit (shape = 3.4, rate = 1.41) (**Table S3**), summarized in **Figure S2**. Quantile-quantile plots showed divergence above a CV of approximately 6, although the empirical and theoretical cumulative distribution functions were stable across the range (**Figure S2**). Based on this model, a probability threshold of 0.05 corresponds to a CV value of 4.8, rounded up to 5.0 (p = 0.0446). This CV threshold was applied in subsequent analyses and produced consistent, biologically reasonable results across multiple samples.

In the final stage of the process, *Owl* score compares the profiled markers against the defined CV threshold. Each haplotype at a marker site is evaluated, while no-called alleles are excluded from the tally. Every called haplotype contributes to the total number of alleles scored. The Score module reports the number of high– and low-CV alleles and summarizes the overall percentage of high-CV alleles as the genome-wide MSI score. In addition to the MSI score, the output includes the percentage of phased sites and the fraction of markers passing quality thresholds.

### Marker filtering

We downloaded genomic DNA sequencing data for 131 samples from HPRC Data Releases 1 and 2 and downsampled to approximately 30-40x depth. Reads were processed with PacBio WGS Variant Pipeline v3.0.2, including alignments to GRCh38_no_alt_analysis_set, variant calls with DeepVariant v1.9.0 and variant phasing and alignment haplotagging with HiPhase v1.5.0. All HPRC sample names used in this study are listed in **Table S2**.

To improve marker reliability, we applied population-level analyses across the HPRC genomes to identify sites that were frequently no-called or consistently scored as MSI-high in our control samples. Among the 164,374 candidate markers, 32,724 (20%) showed at least one no-call, with a median no-call rate of 13%, and 5,872 sites (3.5%) failed in all samples, typically due to insufficient read support. High-CV events were even more common: 122,453 markers (74%) had at least one sample with CV>5, although these events were generally rare at the population scale, with a median of only 2.29% of samples affected per site and just four markers with high CV in all individuals. Using these population-derived metrics, we removed sites where more than 25% (>32) of samples were either no-called or had high CV, yielding 17,790 unreliable markers that were excluded from the final marker set (146,562) distributed with *Owl*.

### Motif specific instability

We quantified motif-level instability by modeling the control cohort using a hierarchical Beta-Binomial. We’ve provided the code and files (**10.5281/zenodo.18437022**). For each sample and motif, we counted the number of unstable events (high) and the total number of evaluated sites (n = high + low). For each motif m, we pooled counts across all 131 HPRC control samples to obtain:

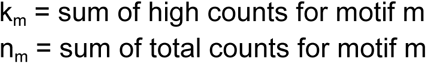

To estimate the underlying instability rate for each motif, we assumed:

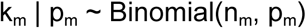

and placed a Beta prior on p_m_ defined by hyperparameters α and β:

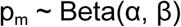

The prior mean was set to the global average instability rate across all motifs in the control cohort, and the prior strength controlled the degree of pooling across motifs. The average instability rate was estimated as μ = 0.0297. The prior strength was set as the median number of evaluated sites across samples, which is given by s = median_m_ (n_m_) = 8. The Beta prior parameters are then set as

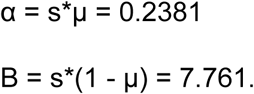

Posterior parameters for each motif were computed using the conjugate update:

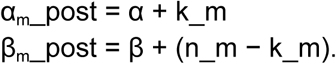

The posterior mean instability rate for motif m was:

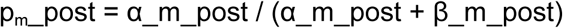

To assess whether a cancer sample showed more instability than expected for a given motif, we used the posterior predictive distribution under the control model. For a new sample with observed high count k_new and total n_new at motif m, the predictive probability of observing k_new or more unstable sites was computed from the Beta-Binomial distribution:

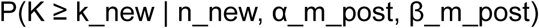

This yields a one-sided posterior-predictive p-value. Motifs whose p-values were small indicate instability greater than expected based on the controls. We applied Bonferroni correction across motifs to control for multiple testing. Under the Bonferroni method, an individual p-value is considered significant if:

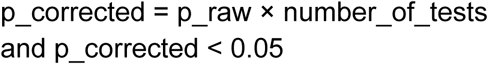

We computed these p-values separately for each cancer sample and retained all motifs with statistically significant excess instability.

### GGAA enrichments

To test for overlap enrichment, we used the R/Bioconductor package regioneR (Gel et al. 2016), running 5,000 permutations with a fixed seed of 42. Analyses were performed on Chr1-22 and ChrX (see **10.5281/zenodo.18437022**).

| sample | condition | p-value | z-score |
| --- | --- | --- | --- |
| TC32 | all-high-motifs | 0.0004997501249 | -4.2704 |
| TC32 | ggaa-high-motifs | 0.0004997501249 | 18.9051 |
| CHLA10 | all-high-motifs | 0.00299850075 | -2.8736 |
| CHLA10 | ggaa-high-motifs | 0.0004997501249 | 17.7504 |
| N/A | ggaa-markers | 0.0004997501249 | 21.6137 |

### DRAGEN MSI analysis

Illumina exome sequencing was performed on an Anaplastic Astrocytoma tumor (caVal-13) and its matched normal using the IDT xGen Exome Research Panel v2.0. Sequencing was carried out to achieve an average coverage of 300x across targeted regions. Sequencing data were processed with the Illumina DRAGEN v4.0.3 to assess MSI in both tumor-normal and the tumor-only mode. For the tumor only mode, a panel of normals with 90 samples was used.

## Acknowledgements

We would like to acknowledge the Human Pangenome Reference Consortium (BioProject ID: PRJNA730823) and its funder, the National Human Genome Research Institute (NHGRI). M.S.F. would like to thank Braden’s Hope for Childhood Cancer, Elizabeth and Monte McDowell, the Black & Veatch Foundation, and Big Slick for their generous support, as well as the Children’s Mercy Research Institute Genomics Core for their assistance with sequencing. PRECISE is a programme of the Consortium for Clinical Research and Innovation, Singapore (CRIS). NPM Phase II is supported by the National Research Foundation, Singapore (NRF) under the RIE2020 White Space (MOH-000588 and MOH-001264) and administered by the Singapore Ministry of Health (MOH) through the National Medical Research Council (NMRC) Office, MOH Holdings Pte Ltd. NPM Phase III is supported by the Singapore MOH through the NMRC Office, MOH Holdings Pte Ltd under the NMRC RIE2025 NPM Phase III Funding Initiative (MOH-001734)

## Author Contributions and conflicts of interest

ZK, KPC, MJPC, ED, and ME contributed to the design of the software. ZK, KPC, BY, and GDSB carried out analyses. ZK, KPC, MJPC, BY, LL, WJR, GDSB, ED, KKH, PT, TP, ERL, SNF, MSF, and MAE wrote the manuscript. WJR, KI, SB, EF, MM, and EO contributed to data generation and or processing of samples. ZK, KPC, ED, WJR, GDSB, and MAE are employees and shareholders of PacBio. ZK is a shareholder of Phase Genomics.

## Data Availability

**Analysis:**

The statistical analysis scripts and associated data files can be found at Zenodo (**10.5281/zenodo.18437022**).

**HPRC**:

HPRC sequencing data can be found at. We used 131 easily accessible and curated samples. https://github.com/human-pangenomics/hprc_intermediate_assembly/tree/main/data_tables/sequencing_data

**CASTLE cell lines**:

CASTLE cell lines data (HCC1954, HCC1937, H1437, H2009, Hs578T) is available through NCBI SRA BioProject PRJNA108684. Accession IDs and sequencing platforms are described on CASTLE’s GitHub page: https://github.com/CASTLE-Panel/castle. For COLO829 and HCC1395, the data can be found on PacBio’s Datasets page: https://downloads.pacbcloud.com/public/revio/2023Q2/HCC1395/ and https://downloads.pacbcloud.com/public/revio/2024Q4/WGS/COLO829.

**The Ewing Sarcoma cell line samples**:

https://www.ncbi.nlm.nih.gov/sra/PRJNA1402985 Singapore: Waiting for Kie-Kyon to reply

**CaVal13 sample**:

The participant did not give written consent for their underlying sequence data to be deposited in a public repository. Specific inquiries about the case can be directed to the corresponding author.

## Supplemental Materials

**Supplemental Figure 1.**
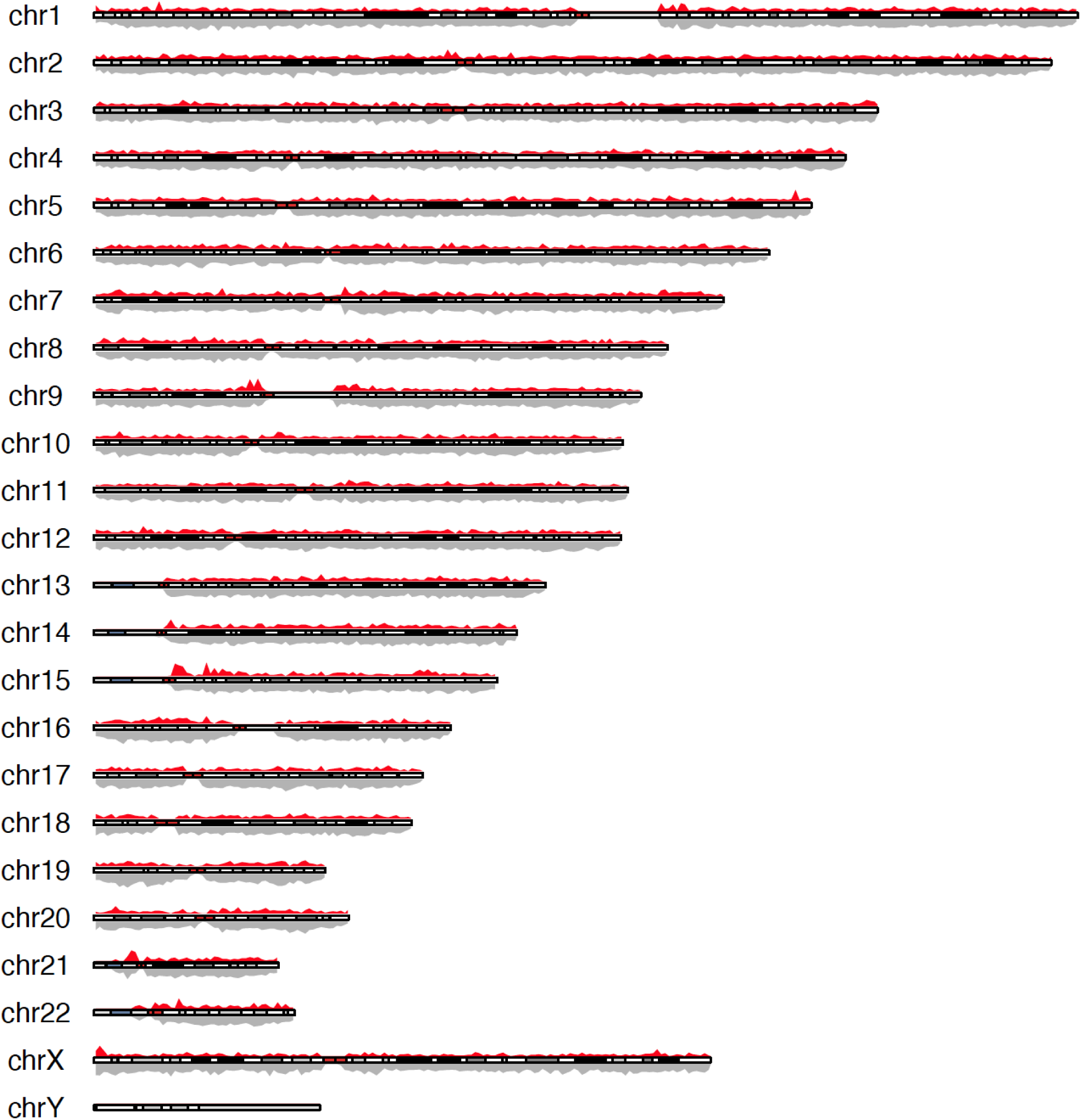
Density of marker sites along GRCh38. The density is smoothed in 1Mb windows. The grey density track is for all the marker sites, and red is for marker sites with extreme variability (CV > 20) in one or more analyzed samples. Chromosome Y was excluded from the *Owl* marker set.

**Supplemental Figure 2.**
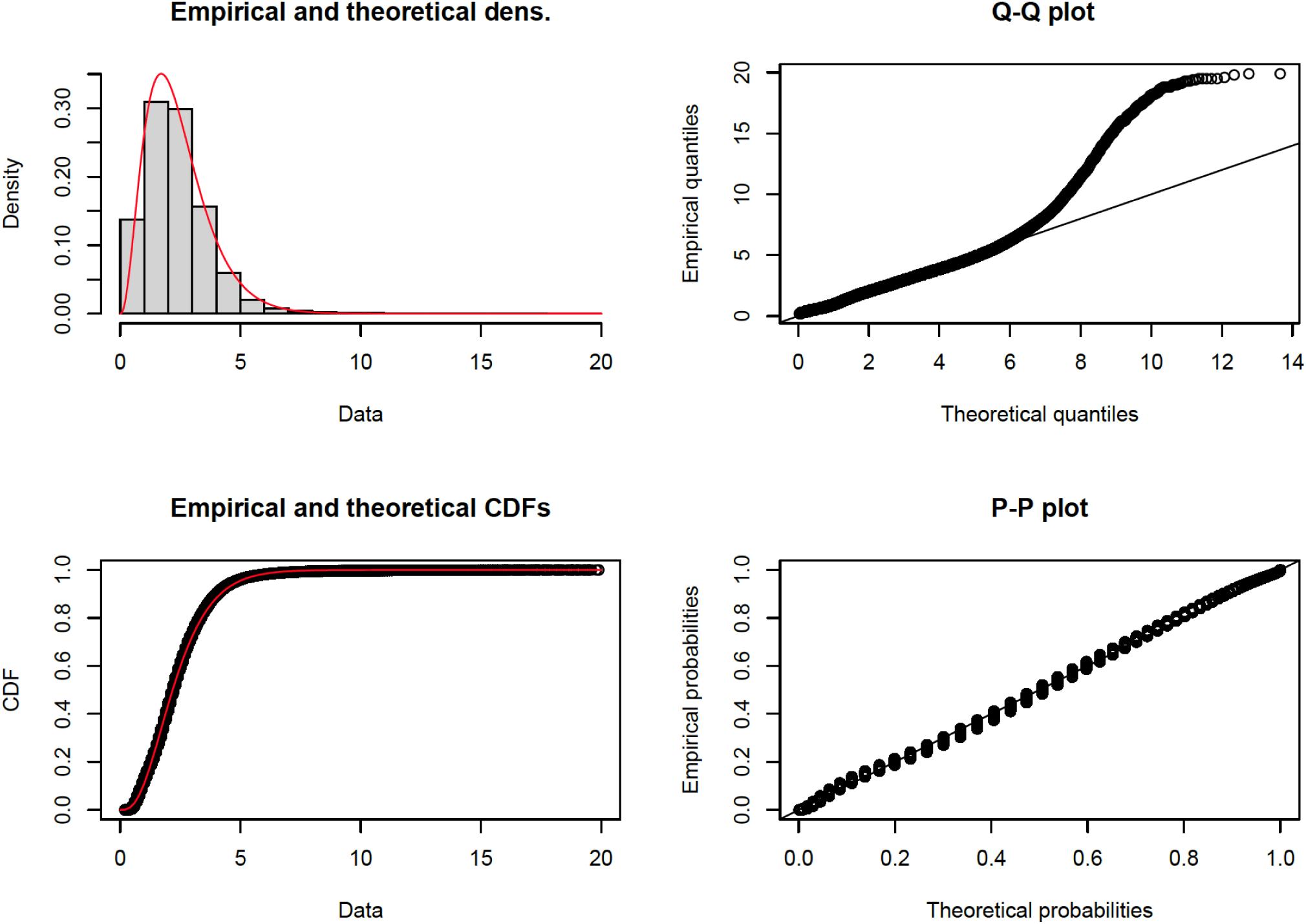
Summary of the Gamma distribution fit to NA12878 CV scores. The data were filtered to exclude zero values (no variation), and extremely high values (CV > 20).

**Supplemental Figure 3.**
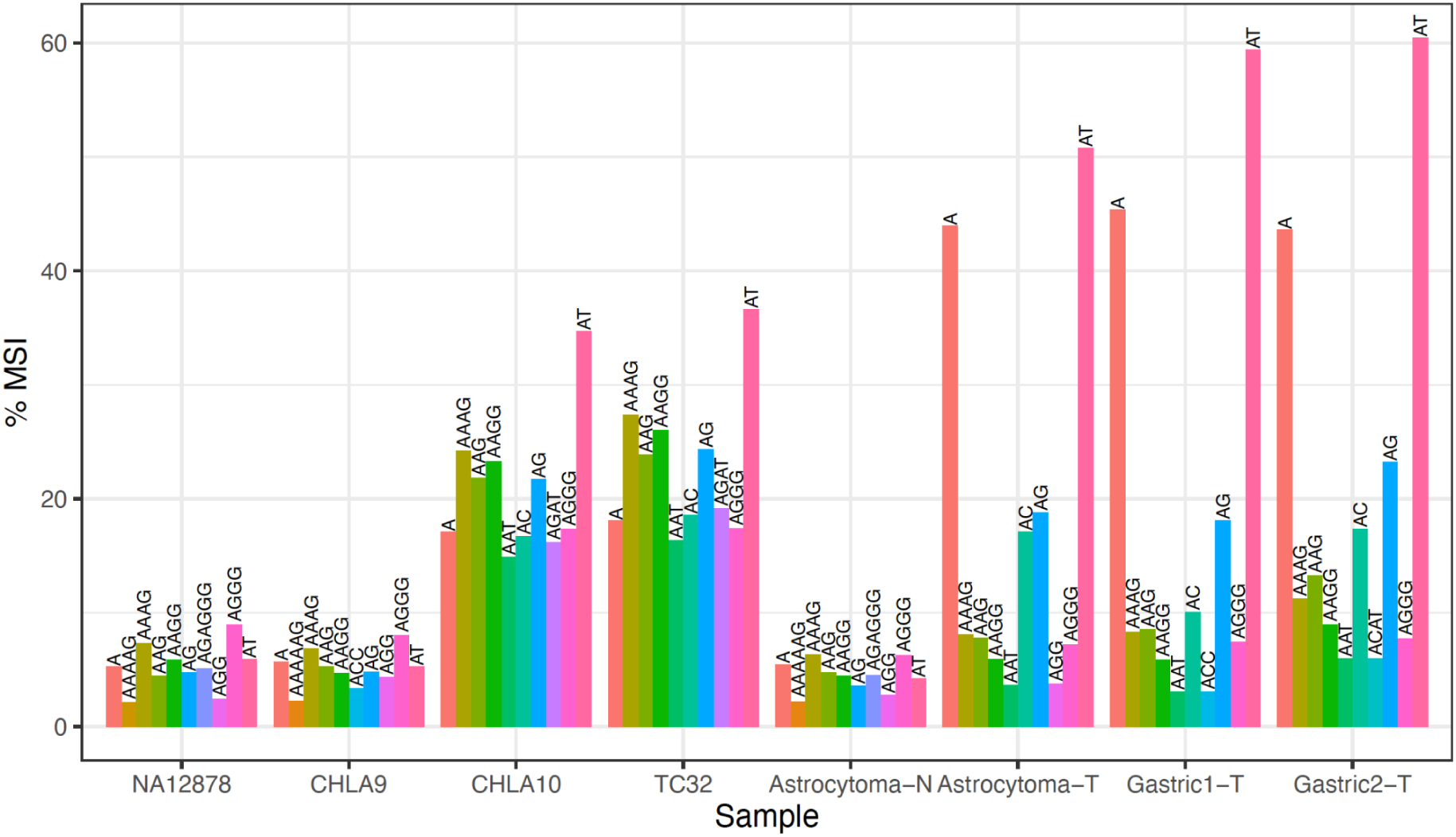
The breakdown of the top-ten highest MSI motifs in select samples. From left to right, NA12878 is a control sample (MSI-stable), CHLA9 is a Ewing sarcoma cell line (MSI-stable), CHLA10 (MSI-high) is an isogenic sample of CHLA9 after chemotherapy, diffuse Astrocytoma control and tumor, and two gastric cancers (MSI-high). The plot shows the top-ten highest scoring motifs, for each sample reported in **Table S4**.

**Supplemental Figure 4.**
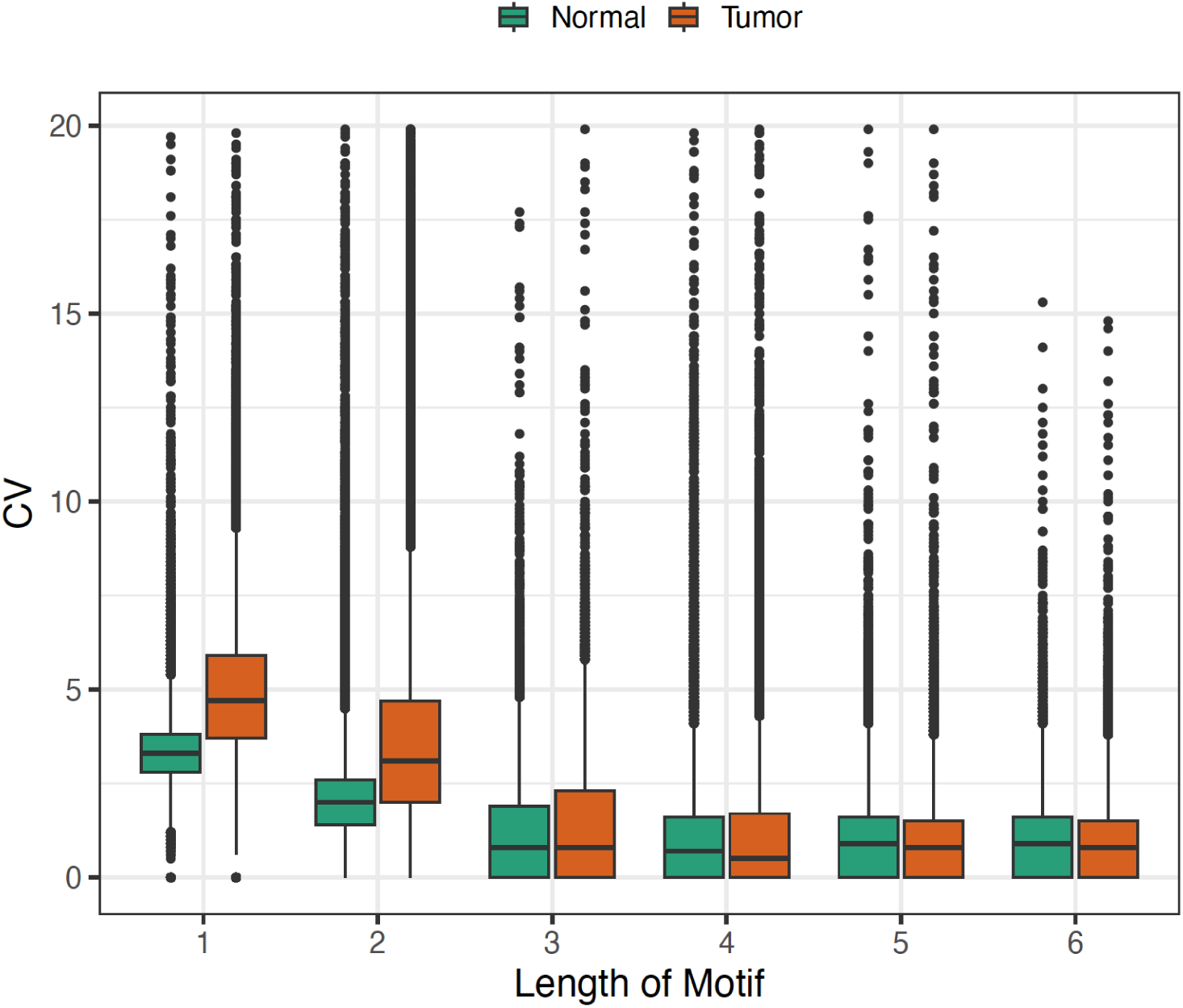
A comparison of the Coefficient of Variance (CV) for the CaVal13 paired Tumor/Normal sample. Values with CV scores > 20 were excluded. Each data represents an alleles CV score. The values are binned by the motif repeat unit length. Most of the CV signals differentiating tumor from normal are homopolymers and di-nucleotides.

**Supplemental Figure 5.**
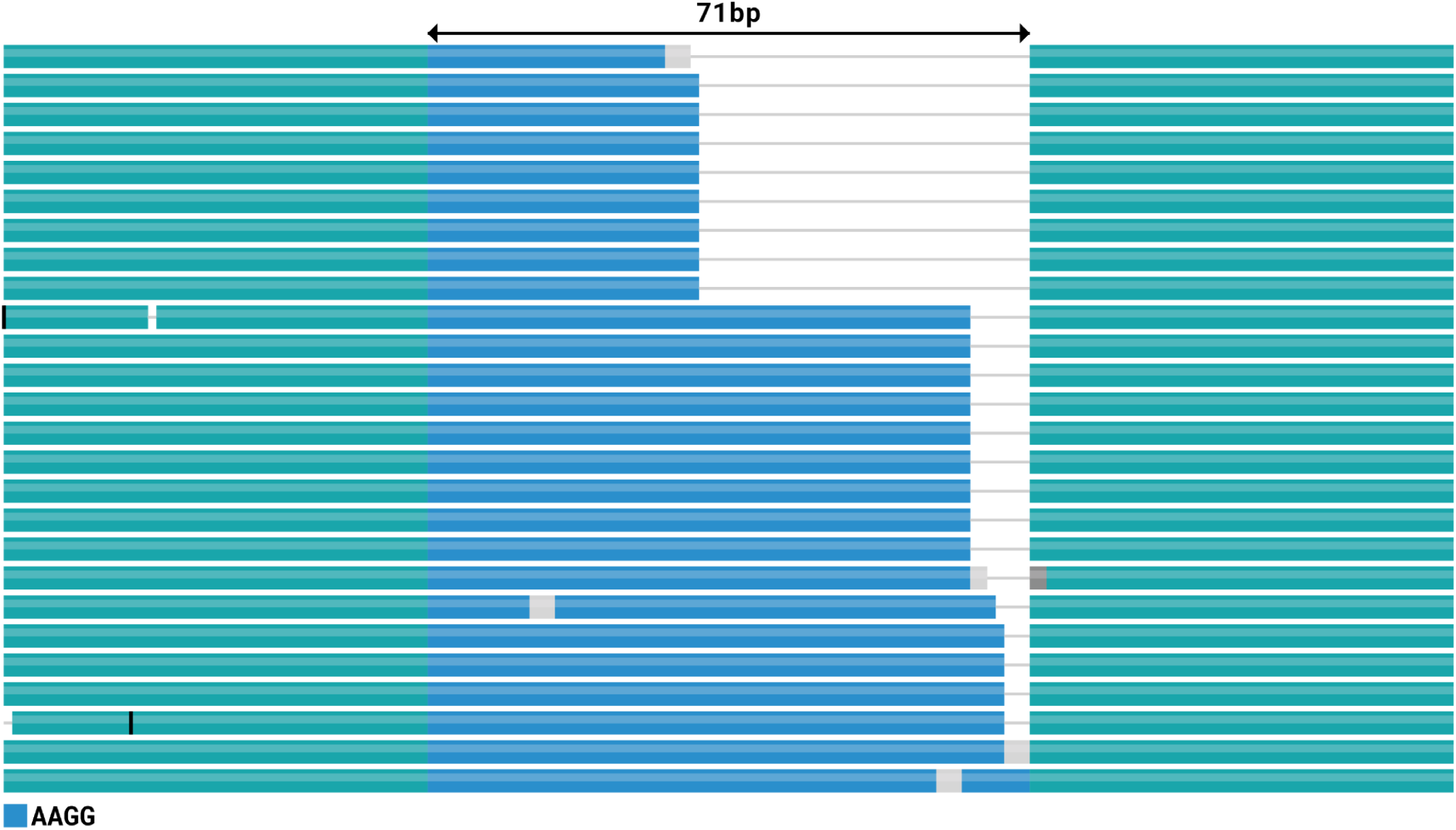
Repeat length distribution plot for TC32 Ewing Sarcoma cell line. The figure shows the repeat lengths for haplotype 2 for the (GGAA is lexicographically minimized to AAGG) chr7:148839053-148839107 marker. The average repeat length (blue) is 58.5, and the CV score is 29. Within a ∼7kb window (chr7:148,836,124-148,842,827) there are several SNVs supporting correct phasing of the reads in haplotype two. The region shown directly overlaps with a distal enhancer-like feature (EH38E2599658) in the third intron of the *EZH2* gene.

